# Wanted not, wasted not: Searching for non-target taxa in environmental DNA metabarcoding by-catch

**DOI:** 10.1101/2021.12.08.471726

**Authors:** Camila Duarte Ritter, Giorgi Dal Pont, Paula Valeska Stika, Aline Horodesky, Nathieli Cozer, Otto Samuel Mader Netto, Caroline Henn, Antonio Ostrensky, Marcio R. Pie

**Affiliations:** Grupo Integrado de Aquicultura e Estudos Ambientais, Departamento de Zootecnia, Universidade Federal do Paraná, Rua dos Funcionários, 1540, Juvevê, 80035-050 Curitiba, PR, Brazil; ATGC Genética Ambiental LTDA. Rua dos Funcionários 1540, Juvevê, Curitiba, PR Brazil, 80035-050; Eukaryotic Microbiology, Faculty of Biology, University of Duisburg-Essen, Universitätsstrasse 5, D-45141 Essen, Germany; Itaipu Binacional. Divisão de Reservatório - MARR.CD, Avenida Tancredo Neves, 6731, Foz do Iguaçu, Paraná, CEP 85866-900, Brazil; Biology Department, Edge Hill University, Ormskirk, Lancashire, L39 4QP, United Kingdom

**Keywords:** Amplicon sequence variants, Birds, Fishes, High throughput sequencing, Mammals, Neotropics, Vertebrata

## Abstract

Metabarcoding of environmental DNA is based on primers that are specific to the target taxa (e.g. bacteria, zooplankton, fishes). However, due to the nature of the commonly used protocols, regardless of the chosen primers, several sequences of non-target species will inevitably be generated, but are usually discarded in commonly used bioinformatics pipelines. These non-target sequences might contain important biological information about the presence of other species in the studied habitats and its potential for ecological studies is still poorly understood. Here, we analyzed the presence of mammal and bird species in aquatic environmental samples that were originally amplified targeting teleost fish species. After all cleaning and checking steps, we kept 21 amplicon sequence variants (ASVs) belonging to mammals and ten to birds. Most ASVs were taxonomic assigned to farm/domestic animals, such as cats, cows, and ducks. Yet, we were able to identify a native semi-aquatic mammal, the capybara, in the samples. Four native bird species and a non-native potentially invasive bird (*Corvus* sp.) were also detected. Although the data derived from these samples for mammals and birds are of limited use for diversity analyses, our results demonstrate the potential of aquatic samples to characterize non-aquatic birds and highlight the presence of a potentially invasive species that had not been recorded before in the region.

## 1. Introduction

In recent decades, environmental DNA (eDNA) metabarcoding revolutionized our ability to efficiently sample and monitor a wide range of taxa (Seymour et al., 2020; Yang et al., 2021). Although eDNA metabarcoding is a cost-effective technique when compared with traditional surveys (Shokralla et al., 2015) or other molecular techniques such as shotgun sequencing (Stat et al., 2017), the cost of amplification and sequencing can pose an important limitation for countries in the Global South. In this context, optimizing the results obtained from eDNA metabarcoding sequencing is highly desirable.

A crucial step of eDNA metabarcoding studies is the choice of genetic marker and primers. The chosen genetic marker should be variable enough to distinguish between target species, whereas the used primers should be specific enough to avoid amplifying non-target taxa (Collins et al., 2019; Leese et al., 2020). However, as any given environmental sample contains a myriad of DNA from entire communities, amplification of non-target taxa is inevitable. Usually, the bioinformatics of taxonomic assignment discards sequences that do not belong to the target taxa (Andújar et al., 2018; Burgess, 2001). Yet, these non-target sequences can provide valuable information about important organisms present in the sample, such as threatened or invasive species, or even seasonal variation in the composition of non-target communities (Mariani et al., 2021).

Aquatic environments have been extensively studied for a wide range of taxa, from viruses to eukaryotic metazoans (Alberti et al., 2017), including many vertebrate taxa (Stat et al., 2017). The most abundant and diverse vertebrates on Earth—the teleost fishes (Osteichthyes) that dominate the aquatic realm — have been widely studied through eDNA metabarcoding in marine and freshwater systems (McElroy et al., 2020), both locally (Sales et al., 2021) and globally (Miya et al., 2020). However, aquatic samples are also able to successfully record other vertebrate classes, such as mammals (Harper et al., 2019; José et al., 2021; Sales et al., 2020).

Although most studies that detected mammals and birds used more general primers for vertebrates (Andruszkiewicz et al., 2017; Closek et al., 2019), a recent study identified such organisms from primers originally designed to amplify teleost fishes (Mariani et al., 2021). Here we explore, for the first time in the Neotropical region, the potential of metabarcoding-derived sequences to identify non-target vertebrates from aquatic environmental samples. We used data from fish monitoring of the Itaipu dam and associated fish pass system in the Paraná River, in South Brazil (Dal Pont et al., 2021). We also discuss some potential caveats and limitations of this approach.

## 2. Material and Methods

Our sampling design is described in Dal Pont et al. (2021). In brief, six sites were sampled in the Piracema fish pass, including a site on the reservoir dam, four in the fish pass and one in the Paraná River in 2019 and three sites that were re-sampled in 2020. The sites were sampled in sextuplicate, each including one liter of water filtered using nitrocellulose membranes (0.45-µm pore). Filters were kept in 100% ethanol under refrigerated conditions. Total DNA from samples and three negative controls were extracted using magnetic beads. We amplified the 12S rRNA gene using the MiFish primers designed by Miya et al. (2015) to yield 163–185 bp fragments targeting teleost fish. The samples and the three negative controls were sequenced with Illumina MiSeq (Illumina, USA). The raw sequences are deposited in GenBank under Bioproject PRJNA750895 (biosamples SAMN20500524 – SAMN20500577).

To determine amplicon sequence variants (ASVs), we first removed primers with the Cutadapt package (Martin, 2011) in Python v.3.3 (Van Rossum and Drake, 2009), and then used the DADA2 package (Callahan et al., 2016) in R v. 4.0.2 (R Core Team, 2021) to quality filter reads, merge sequences, remove chimeras, and to infer ASVs. ASVs present with a proportion > 0.01% of reads across all three negative controls were discarded.

For taxonomic inference, we build a reference dataset of 12S mitochondrial DNA sequences for fish, mammal, and bird taxa that have been historically recorded in the Itaipu area using the available data in GenBank (Benson et al., 2018). A total of 75 bird, 126 fish, and 78 mammal species had sequences available and were used. For fishes, we added an in-house database which included sequences for 42 additional species (Dal Pont et al., 2021). Finally, we blasted the obtained ASVs sequences with our reference database to verify the taxonomic composition using the “Blastn” function of the program Blast+ (Camacho et al., 2009) for the 10 best hits and an e-value of < 0.001. We kept ASVs that matched a species from our reference at minimum level of 75% similarity. Inconsistent results were checked manually. ASVs blasted > 98% similarity was considered the matched species. All other ASVs were blasted in GenBank as an additional check and then replaced if there was a match with a superior e-value. ASVs with similarity between 96 to 98% were considered in the same genus, 90 to 96 the same family. For similarities between 75 to 90 the ASVs were just considered for the class. We used the metagMisc v. 0.0.4 (Mikryukov, 2019), phyloseq v. 1.36.0 (McMurdie and Holmes, 2013), and tidyverse v. 1.3.0 (Wickham, 2017) packages for data curation and ggplot2 v. 3.3.2 (Wickham, 2016), plotly v. 4.10.0 (Sievert, 2020), and patchwork v.1.1.1 (Pedersen, 2019), for data visualization in R v. 4.1.1 (R Core Team, 2021). The script is available as Appendix 1.

## 3. Results

We obtained a total of 17,616,032 reads of raw sequence. After all cleaning steps, we retained a total of 2,280,447 reads belonging to 7,096 ASVs. After we removed ASVs with a proportion of > 0.1% of reads present in the sum of three negative controls and taxonomically assigned the ASVs, we kept 994,251 (44% of total) sequences belonging to 220 ASVs with at least 75% similarity of one species in our reference database. As expected, most ASVs (189 ASVs in a total of 966,610 reads) belong to fishes, followed by mammals (26,127 reads belonging to 21 ASVs), and only 10 ASVs (1,514 reads) assigned as birds (Fig. 1).

**Figure 1.**
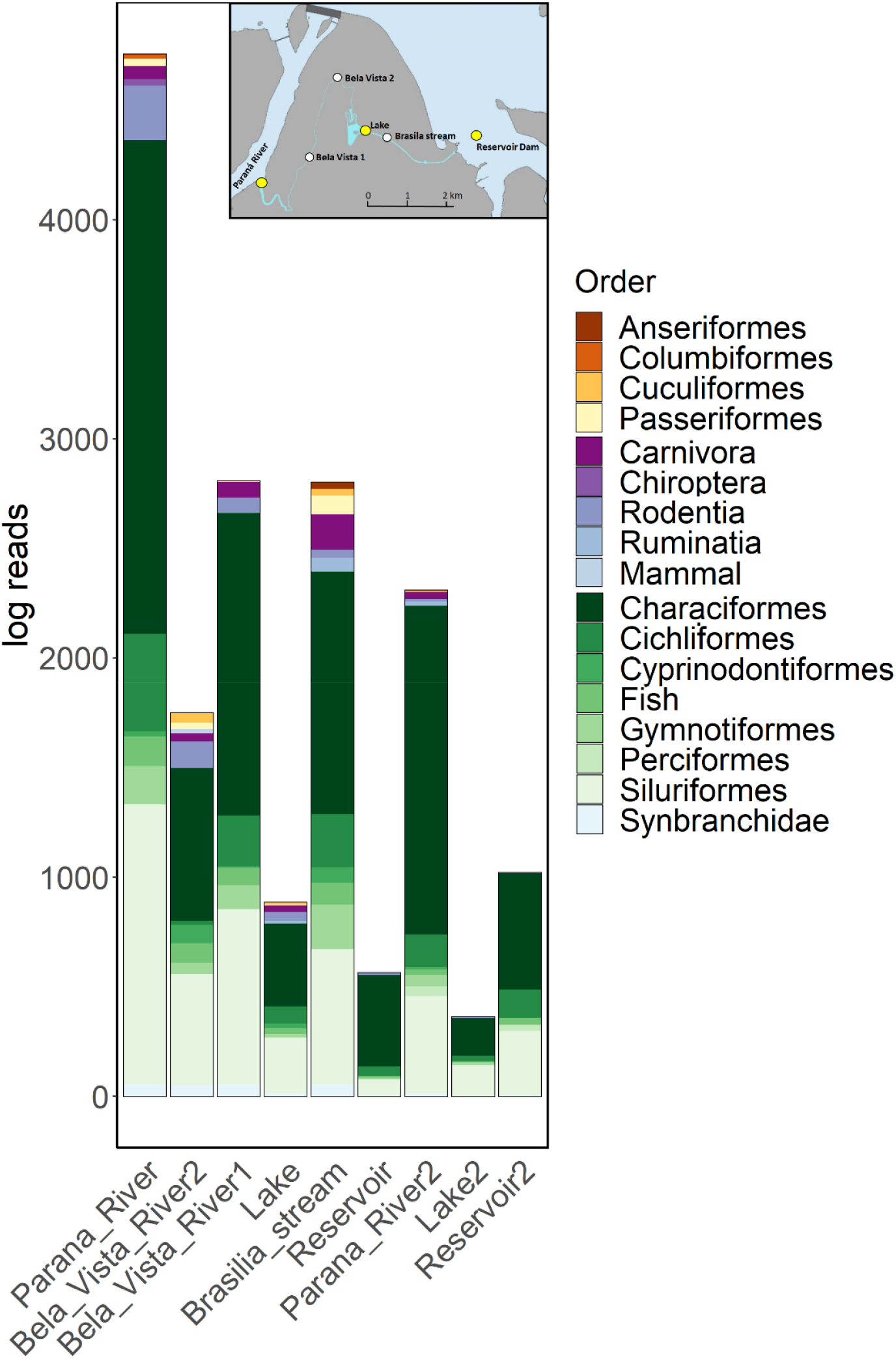
Taxonomic composition by order and class in Piracema fish pass. Inset panel show the point location from left to right as the bars order sampled in 2019. Point in yellow were re-sampled in 2020 (last three bars).

From the 21 ASVs assigned as mammal, 12 ASVs were assigned at species level (five species including dog, cat, mouse, and cow), one at genus level, seven at family level, and one only as a mammal. From the ten ASVs assigned as birds, seven were assigned at species level (six species, including a *Corvus sp*.), one at genus level, and two at family level (Table S1). Most non-fish ASV were recorded in 2019, with higher abundance in the Parana River and the higher number of ASVs in the Brasilia stream locality (Fig. 2).

**Figure 2.**
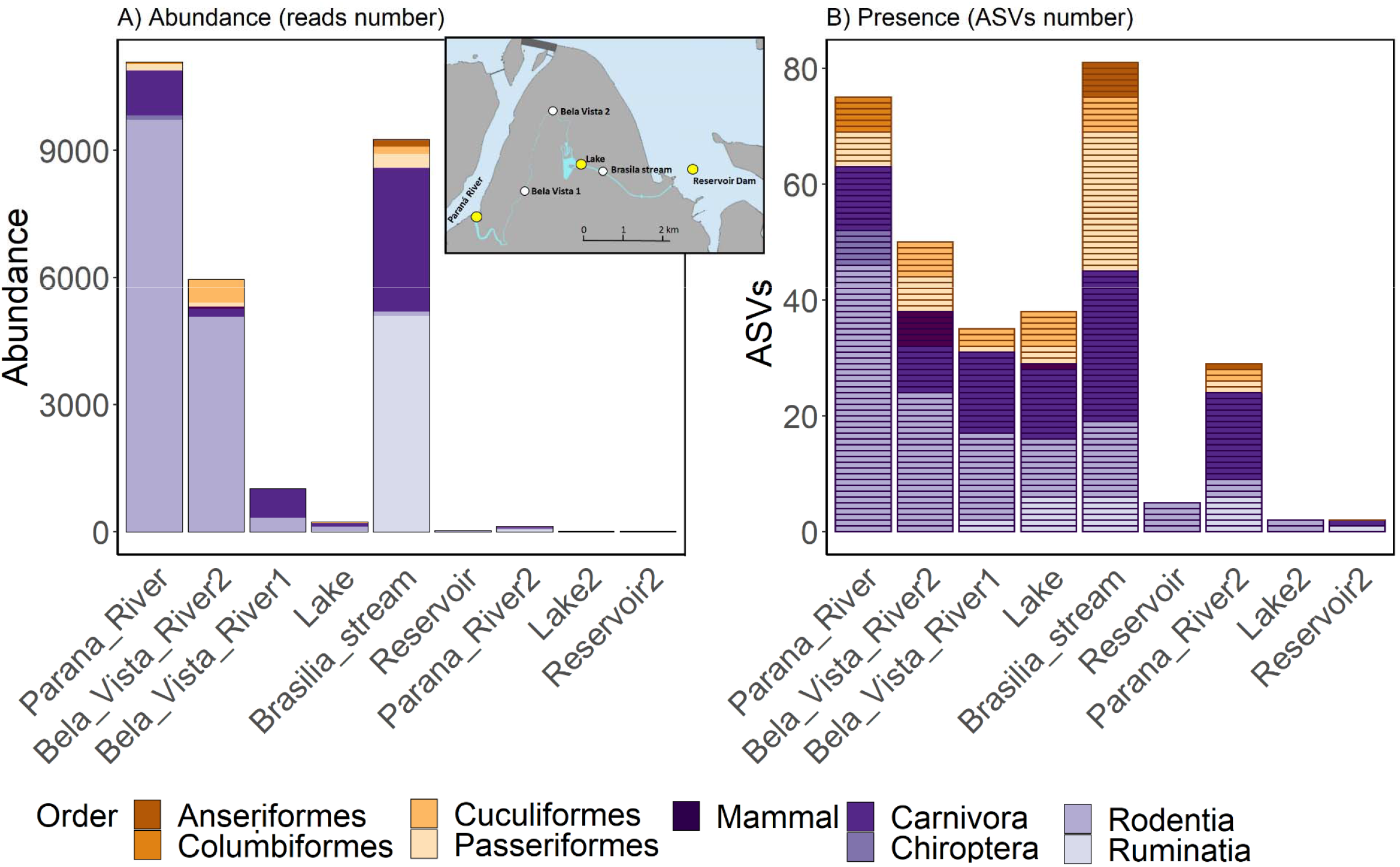
Taxonomic composition order abundance (A) and ASVs number (B) for birds and mammals registered in the Piracema fish pass. Inset panel show the point location from left to right as the bars order sampled in 2019. Point in yelow were re-sampled in 2020 (last three bars). One ASV in Bela Vista River 2 point assigned as mammal could not been assigned by any order by the low similarity (79.6%) and was kept as “mammal”.

## 4. Discussion

Our results demonstrate that environmental DNA metabarcoding by-catch is a viable source of genetic information for non-target species. Indeed, this is the second study to explore this possibility (Mariani et al., 2021), and the first in the Neotropics. Interestingly, contrary to that study, our data recorded mostly farm/urban animals, such as cat, dog, rat, cow, and duck. It is interesting to note that these records were detected in samples from 2019, whereas in 2020 almost no sequences of non-target organisms were found. In 2020 there was an extreme drought in southeastern Brazil (de Oliveira Bueno et al., 2020), which potentially impacted fish assemblages (Dal Pont et al., 2021) and probably other organisms present in the region as well. Another relevant result is the lower number ASVs recorded in the reservoir site, probably associated with the high water volume near the Itaipu Hydroelectric Power Plant dam and, then low eDNA density.

A unique native non-farm / urban mammal species recorded was the capybara *Hydrochoerus hydrochaeris*. Capybara is the largest living rodent of the world (Nowak and Walker, 1999) with a semi-aquatic habit (Corriale and Herrera, 2014) and occurring throughout most of South America (Moreira et al., 2013), being common in the region (Corriale and Herrera, 2014; Dias et al., 2020). Other mammals were only assigned at the family level, including other rodents (Cricetidae and Hydrocharidae), one bat (Phyllostomidae), and one carnivore (Procyonidae), that is a New World family (Duszynski et al., 2018). Although two species of Procyonidae, *Nasua nasua* and *Procyon crancrivorous*, are common in the region (Brocardo et al., 2019), these species were present in our reference database matching with a low similarity. These sequences may have low similarity due to the lack of publicly available representative sequences of the species which them belong, highlighting the need of further studies in the Neotropical region.

For birds, beyond the very common non-native species house sparrow (*Passer domesticus*) and duck (*Cairina moschata*), we recorded four additional species. From these, three are native and common species, demonstrating the potential of eDNA of aquatic samples to record non-aquatic bird species. One species we found was a crow (*Corvus* sp.) that is non-native of South America (Burton and Burton, 2002). The corvid species occurring in the region is *Cyanocharax chrysops* (http://www.aultimaarcadenoe.com.br/aves-do-pq-nacional-do-iguacu/), and although no sequence for this species is available other species of the genera had it and match with very low similarity, being the *Corvus* sp. the most probable species. To the best of our knowledge, only one species of crow is known to occur in Brazil, *Corvus albus*, which was reported for the first time in 2004 (Silva e Silva and Olmos, 2007), followed by additional records (Adelino et al., 2017; Lima and Kamada, 2009). This species is considered a native invader in its home range that has benefited from human infrastructure (Cunningham et al., 2016) and is a potentially invasive species in Brazil (Adelino et al., 2017). It is alarming, since invasive species are detrimental to both biodiversity, ecosystem process, human welfare, and economy (Blackburn et al., 2014). In particular, *C. albus* has established populations outside their native range in several countries, where they are responsible for ecological impacts (Ryall, 1992), economic loss (Kamel, 2014), and human health problems (Yap and Sodhi, 2004). Although the record derived from metabarcoding data targeting fishes should be considered carefully, it indicates a possible occurrence of a potentially invasive species that deserves to be further investigated.

## 5. Conclusions

Metabarcoding studies generate hundreds to thousands of non-target sequences. Although this data is severing biased, it can contain important biological information. It is important to note that it is unlikely that environmental DNA metabarcoding by-catch will provide sufficient information for comprehensive surveys and diversity estimates of non-target species. However, we envision two particularly useful applications of this approach. First, it might provide valuable information on population fluctuations of the most common species that live close or associated to bodies of water, such as the capybara. Second, it might show general trends in the anthropic influence in the region, as evidence of native species might be slowly replaced by domestic ones.

In addition, we provide evidence for the potential presence on an invasive species that may be controlled before it becomes a major problem. Also, we showed that aquatic samples are suitable to detect Neotropical bird species. The use of primers specific for birds in aquatic samples can optimize sampling in highly diverse and remote areas.

## Acknowledgements

Funding was provided by Itaipu Binacional to AON (grant #4500049847). We thank Itaipu Binacional for providing postdoctoral fellow grants to GDP and NC and the Alexander van Humboldt foundation for providing a postdoctoral fellow grant to CDR. We also thank Conselho Nacional de Desenvolvimento Científico e Tecnológico (CNPq) for awarding AO and MRP with research fellowship grants (grants #304633/2017-8 and #302904/2020-4, respectively).

## Author Contribution Statement

M.P. study conceptualization, G.D.P, A.H, N.C., O.S.M.N., A.O., MP. experimental design. A.O.A, P.V.S., and E.B. molecular analysis. C.D.R. bioinformatics and statistical analysis. A.O. an M.P. fund-raising. C.D.R. wrote the first version with the collaboration of all authors. All authors read and approved the final version.

