## Appendix1 for "Wanted not, wasted not: Searching for non-target taxa in environmental DNA metabarcoding by-catch": Appendix1.html

Itaipu metabarcoding bycatch paper


### Itaipu metabarcoding bycatch paper

###### Camila Duarte Ritter

###### November 2021

### Setup

```
knitr::opts_chunk$set(echo = F)

#Library needed packages

library(tidyverse)
```

```
## -- Attaching packages --------------------------------------- tidyverse 1.3.1 --
```

```
## v ggplot2 3.3.5     v purrr   0.3.4
## v tibble  3.1.5     v dplyr   1.0.7
## v tidyr   1.1.4     v stringr 1.4.0
## v readr   2.0.2     v forcats 0.5.1
```

```
## -- Conflicts ------------------------------------------ tidyverse_conflicts() --
## x dplyr::filter() masks stats::filter()
## x dplyr::lag()    masks stats::lag()
```

```
library(phyloseq)
library(metagMisc)
```

```
## 
## Attaching package: 'metagMisc'
```

```
## The following object is masked from 'package:purrr':
## 
##     some
```

```
library(patchwork)
library(plotly)
```

```
## 
## Attaching package: 'plotly'
```

```
## The following object is masked from 'package:ggplot2':
## 
##     last_plot
```

```
## The following object is masked from 'package:stats':
## 
##     filter
```

```
## The following object is masked from 'package:graphics':
## 
##     layout
```

### My Data

```
#Read in the relevant files:

#metadata
metadata <- read.csv("metadata_Itaipu2.csv")
row.names(metadata) <- metadata$Sample#here we name the rows
head(metadata)
```

```
##      Sample     Group     Local     Year  Symbol
## P1.1   P1.1 Reservoir Reservoir Nineteen Diamond
## P1.2   P1.2 Reservoir Reservoir Nineteen Diamond
## P1.3   P1.3 Reservoir Reservoir Nineteen Diamond
## P1.4   P1.4 Reservoir Reservoir Nineteen Diamond
## P1.5   P1.5 Reservoir Reservoir Nineteen Diamond
## P1.6   P1.6 Reservoir Reservoir Nineteen Diamond
```

```
#data at ASV level
ASVs <- read.csv("ASVsl.csv")
#head(ASVs)

#### transform table to matrix

#ASVs
ASVs_M <- as.matrix(ASVs[, -1])#here we remove the name column
dimnames(ASVs_M) <- list(ASVs[,1], colnames(ASVs)[-1])#here we name the rows
```

#### Construct phyloseq objects

```
# ASV
tax <- read.csv("tax.csv")
tax_M <- as.matrix(tax)  #here we remove the name column
row.names(tax_M) <- tax$ASV_ID  #here we name the rows
head(tax_M)
```

```
##         ASV_ID    Domain Phylum Class  Order           Family       
## ASV1    "ASV1"    "Fish" "Fish" "Fish" "Characiformes" "Characidae" 
## ASV100  "ASV100"  "Fish" "Fish" "Fish" "Characiformes" "Crenuchidae"
## ASV1017 "ASV1017" "Bird" "Bird" "Bird" "Anseriformes"  "Anatidae"   
## ASV102  "ASV102"  "Fish" "Fish" "Fish" "Characiformes" "Curimatidae"
## ASV1024 "ASV1024" "Fish" "Fish" "Fish" "Gymnotiformes" "Apteronidae"
## ASV1026 "ASV1026" "Fish" "Fish" "Fish" "Characiformes" "Anostomidae"
##         Genus           Species                   
## ASV1    "Astyanax"      "Astyanax altiparanae"    
## ASV100  ""              ""                        
## ASV1017 "Cairina"       "Cairina moschata"        
## ASV102  ""              ""                        
## ASV1024 "Apteronotus"   ""                        
## ASV1026 "Megaleporinus" "Megaleporinus obtusidens"
```

```
AbuASV <- phyloseq(otu_table(ASVs_M, taxa_are_rows = TRUE), tax_table(tax_M), sample_data(metadata))
```

### Order distribution with fishs included

```
newSOrder = c("Parana_River", "Bela_Vista_River2", "Bela_Vista_River1", "Lake", "Brasilia_stream",
    "Reservoir", "Parana_River2", "Lake2", "Reservoir2")

taxborder <- c("#66000000", "#66000000", "#66000000")
colTrans <- rep("#66000000", 236)


taxCols <- c("#bfd3e6", "#993404", "#ffeda0", "#810f7c", "#00441b", "#8856a7", "#238b45",
    "#d95f0e", "#fec44f", "#41ab5d", "#74c476", "#a1d99b", "#bfd3e6", "#fff7bc",
    "#c7e9c0", "#8c96c6", "#9ebcda", "#e5f5e0", "#e5f5f9")

# Logarithmic transformation as in Anderson et al., 2006
logAbu <- phyloseq_standardize_otu_abundance(AbuASV, method = "log")

fig_comp <- plot_bar(logAbu, x = "Group", fill = "Domain") + geom_bar(aes(color = Domain,
    fill = Order), stat = "identity", position = "stack") + scale_color_manual(values = taxborder,
    0.2, "Order") + scale_fill_manual(values = taxCols, "Order") + ylab("log reads") +
    theme(axis.text = element_text(size = 20), axis.title.x = element_blank(), axis.title.y = element_text(size = 22),
        strip.text = element_text(size = 22), legend.title = element_text(size = 20),
        legend.text = element_text(size = 20), axis.ticks.x = element_blank(), plot.title = element_text(size = 15L),
        plot.caption = element_text(size = 12L), axis.text.x = element_text(angle = 45,
            hjust = 1), panel.grid.major.x = element_blank(), panel.grid.major.y = element_blank(),
        panel.grid.minor = element_blank(), panel.background = element_blank(), axis.line = element_line(colour = "black"),
        panel.border = element_rect(colour = "black", fill = NA, size = 1)) + guides(color = "none")
```

```
## Warning in psmelt(physeq): The sample variables: 
## Sample
##  have been renamed to: 
## sample_Sample
## to avoid conflicts with special phyloseq plot attribute names.
```

```
fig_comp$data$Group <- as.character(fig_comp$data$Group)
fig_comp$data$Group <- factor(fig_comp$data$Group, levels = newSOrder)

ordertax <- c("Bird", "Mammal", "Fish")
fig_comp$data$Domain <- as.character(fig_comp$data$Domain)
fig_comp$data$Domain <- factor(fig_comp$data$Domain, levels = ordertax)

ggsave("fig_comp2.tiff", fig_comp, width = 8, height = 12, device = tiff)
plotly::ggplotly(fig_comp)
```

### Birds and Mammal order distribution

```
Tetra = subset_taxa(AbuASV, Class != "Fish")

colstetra <- c("#4d004b", "#b35806", "#7f3b08", "#542788", "#8073ac", "#e08214",
    "#fdb863", "#2d004b", "#fee0b6", "#b2abd2", "#d8daeb")

taxborder2 <- c("#7f3b08", "#2d004b")

theme2 <- theme(axis.text = element_text(size = 20), axis.title.x = element_blank(),
    axis.title.y = element_text(size = 22), strip.text = element_text(size = 22),
    legend.title = element_text(size = 20), legend.text = element_text(size = 20),
    axis.ticks.x = element_blank(), plot.title = element_text(size = 15L), plot.caption = element_text(size = 12L),
    axis.text.x = element_text(angle = 45, hjust = 1), panel.grid.major.x = element_blank(),
    panel.grid.major.y = element_blank(), panel.grid.minor = element_blank(), panel.background = element_blank(),
    axis.line = element_line(colour = "black"), panel.border = element_rect(colour = "black",
        fill = NA, size = 1))

fig_comp_tetra_A <- plot_bar(Tetra, x = "Group", fill = "Class") + geom_bar(aes(color = Class,
    fill = Order), stat = "identity", position = "stack") + ggtitle("A) Abundance (reads number)") +
    scale_color_manual(values = taxborder) + scale_fill_manual(values = colstetra,
    "Order") + guides(color = "none") + theme2
```

```
## Warning in psmelt(physeq): The sample variables: 
## Sample
##  have been renamed to: 
## sample_Sample
## to avoid conflicts with special phyloseq plot attribute names.
```

```
presab <- phyloseq_standardize_otu_abundance(Tetra, method = "pa")  #make presence/absence matrix

fig_comp_tetra_B <- plot_bar(presab, x = "Group", fill = "Class") + geom_bar(aes(color = Class,
    fill = Order), stat = "identity", position = "stack") + ggtitle("B) Presence (ASVs number)") +
    scale_color_manual(values = taxborder2) + scale_fill_manual(values = colstetra,
    "Order") + ylab("ASVs") + guides(color = "none") + theme2
```

```
## Warning in psmelt(physeq): The sample variables: 
## Sample
##  have been renamed to: 
## sample_Sample
## to avoid conflicts with special phyloseq plot attribute names.
```

```
fig_comp_tetra_A$data$Group <- as.character(fig_comp_tetra_A$data$Group)
fig_comp_tetra_A$data$Group <- factor(fig_comp_tetra_A$data$Group, levels = newSOrder)

fig_comp_tetra_B$data$Group <- as.character(fig_comp_tetra_B$data$Group)
fig_comp_tetra_B$data$Group <- factor(fig_comp_tetra_B$data$Group, levels = newSOrder)

figCompTetra <- fig_comp_tetra_A + fig_comp_tetra_B + plot_layout(guides = "collect") &
    theme(legend.position = "bottom")
ggsave("figCompTetra.tiff", figCompTetra, width = 12, height = 8, device = tiff)
plotly::ggplotly(fig_comp_tetra_A)
```

```
plotly::ggplotly(fig_comp_tetra_B)
```

### Plot network by Bray-Curtis distance

```
# Make network with Bray Curtis Distance
ig <- make_network(Tetra, "samples", distance = "bray", max.dist = 0.95)
```

```
## Warning in vegdist(structure(c(0L, 0L, 0L, 0L, 0L, 0L, 0L, 0L, 0L, 0L, 0L, : you
## have empty rows: their dissimilarities may be meaningless in method "bray"
```

```
## Warning in vegdist(structure(c(0L, 0L, 0L, 0L, 0L, 0L, 0L, 0L, 0L, 0L, 0L, :
## missing values in results
```

```
cols <- c("#a6cee3", "#cab2d6", "#ffff99", "#fb9a99", "#636363", "#b2df8a", "#1f78b4",
    "#e31a1c", "#33a02c")

forms <- c(15, 16, 16, 16, 16, 17, 0, 1, 2)

set.seed(2)
netPlot <- plot_network(ig, Tetra, type = "samples", point_size = 8, label = NULL,
    color = "Group", shape = "Group", line_alpha = 0.08) + scale_color_manual(values = cols,
    "Locality") + scale_shape_manual(values = forms, "Locality") + theme(axis.text = element_text(size = 20),
    axis.title = element_text(size = 22), strip.text = element_text(size = 22), legend.title = element_text(size = 20),
    legend.text = element_text(size = 20), axis.ticks.x = element_blank(), plot.title = element_text(size = 15L),
    plot.caption = element_text(size = 12L), panel.grid.major.x = element_blank(),
    panel.grid.major.y = element_blank(), panel.grid.minor = element_blank(), panel.background = element_blank(),
    axis.line = element_line(colour = "black"), panel.border = element_rect(colour = "black",
        fill = NA, size = 1), legend.position = c(0.87, 0.25), legend.background = element_rect(fill = "transparent",
        color = "black"))


netPlot$data$Group <- as.character(netPlot$data$Group)
netPlot$data$Group <- factor(netPlot$data$Group, levels = newSOrder)

ggsave("netPlot.tiff", netPlot, width = 12, height = 8, device = tiff)
plotly::ggplotly(netPlot)
```

---


Session Info

```
## [1] "2021-11-26 18:46:29 CET"
```

```
## R version 4.1.1 (2021-08-10)
## Platform: x86_64-w64-mingw32/x64 (64-bit)
## Running under: Windows 10 x64 (build 19044)
## 
## Matrix products: default
## 
## locale:
## [1] LC_COLLATE=English_United States.1252 
## [2] LC_CTYPE=English_United States.1252   
## [3] LC_MONETARY=English_United States.1252
## [4] LC_NUMERIC=C                          
## [5] LC_TIME=English_United States.1252    
## 
## attached base packages:
## [1] stats     graphics  grDevices utils     datasets  methods   base     
## 
## other attached packages:
##  [1] plotly_4.10.0   patchwork_1.1.1 metagMisc_0.0.4 phyloseq_1.36.0
##  [5] forcats_0.5.1   stringr_1.4.0   dplyr_1.0.7     purrr_0.3.4    
##  [9] readr_2.0.2     tidyr_1.1.4     tibble_3.1.5    ggplot2_3.3.5  
## [13] tidyverse_1.3.1
## 
## loaded via a namespace (and not attached):
##  [1] nlme_3.1-152           bitops_1.0-7           fs_1.5.0              
##  [4] lubridate_1.8.0        httr_1.4.2             GenomeInfoDb_1.28.4   
##  [7] tools_4.1.1            backports_1.3.0        bslib_0.3.1           
## [10] vegan_2.5-7            utf8_1.2.2             R6_2.5.1              
## [13] lazyeval_0.2.2         mgcv_1.8-36            DBI_1.1.1             
## [16] BiocGenerics_0.38.0    colorspace_2.0-2       permute_0.9-5         
## [19] rhdf5filters_1.4.0     ade4_1.7-18            withr_2.4.2           
## [22] tidyselect_1.1.1       compiler_4.1.1         cli_3.0.1             
## [25] rvest_1.0.2            Biobase_2.52.0         formatR_1.11          
## [28] xml2_1.3.2             labeling_0.4.2         sass_0.4.0            
## [31] scales_1.1.1           digest_0.6.28          rmarkdown_2.11        
## [34] XVector_0.32.0         pkgconfig_2.0.3        htmltools_0.5.2       
## [37] dbplyr_2.1.1           fastmap_1.1.0          htmlwidgets_1.5.4     
## [40] rlang_0.4.11           readxl_1.3.1           rstudioapi_0.13       
## [43] farver_2.1.0           jquerylib_0.1.4        generics_0.1.1        
## [46] jsonlite_1.7.2         crosstalk_1.2.0        RCurl_1.98-1.5        
## [49] magrittr_2.0.1         GenomeInfoDbData_1.2.6 biomformat_1.20.0     
## [52] Matrix_1.3-4           Rcpp_1.0.7             munsell_0.5.0         
## [55] S4Vectors_0.30.2       Rhdf5lib_1.14.2        fansi_0.5.0           
## [58] ape_5.5                lifecycle_1.0.1        stringi_1.7.5         
## [61] yaml_2.2.1             MASS_7.3-54            zlibbioc_1.38.0       
## [64] rhdf5_2.36.0           plyr_1.8.6             grid_4.1.1            
## [67] parallel_4.1.1         crayon_1.4.2           lattice_0.20-44       
## [70] splines_4.1.1          Biostrings_2.60.2      haven_2.4.3           
## [73] multtest_2.48.0        hms_1.1.1              knitr_1.36            
## [76] pillar_1.6.4           igraph_1.2.7           reshape2_1.4.4        
## [79] codetools_0.2-18       stats4_4.1.1           reprex_2.0.1          
## [82] glue_1.4.2             evaluate_0.14          data.table_1.14.2     
## [85] modelr_0.1.8           vctrs_0.3.8            tzdb_0.1.2            
## [88] foreach_1.5.1          cellranger_1.1.0       gtable_0.3.0          
## [91] assertthat_0.2.1       xfun_0.26              broom_0.7.10          
## [94] viridisLite_0.4.0      survival_3.2-11        iterators_1.0.13      
## [97] IRanges_2.26.0         cluster_2.1.2          ellipsis_0.3.2
```
